# Irinotecan-induced mucositis is associated with variation in the expression of regulatory compounds associated with goblet cells

**DOI:** 10.1101/2025.01.05.631381

**Authors:** Daniel W Thorpe, Bronwen Mayo, John Bresland, Andrea Stringer

## Abstract

Alimentary mucositis (AM) is a common side effect of antineoplastic treatment and a key reason for cessation of treatment, compromising the chance of remission. Gastrointestinal mucin secretion is associated with regulatory compounds nitric oxide and its respective synthases inducible, epithelial and neural nitric oxide synthase (iNOS, eNOS and nNOS, respectively), vasoactive intestinal polypeptide (VIP), and prostaglandin E2 (PGE2). Changes in secretion during mucositis have been demonstrated in numerous studies, however no secretory regulatory signals have been associated in mucositis induced secretory change. The aim of this study is to investigate regulator factors in the gastrointestinal tract involved in mucin secretion, VIP, NOS, and PGE2, which are suspected of being involved in mucositis, specifically through alteration of neural or goblet cells. Tumour-bearing Dark Agouti rats received a single dose of 175 mg/kg of irinotecan (i.p.) and 0.01mg/kg atropine. (s.c.). Rats were killed post treatment at 6, 24, 48, 72, 96 and 120 hours. Samples were collected, immunohistochemistry and real time PCR were performed to analyse the expression of iNOS, eNOS, nNOS, VIP, and PGE2. Following irinotecan treatment, staining intensity for iNOS positive goblet cells in the crypts of the jejunum decreased significantly at 48 h (p < 0.05). VIP, and PGE2 positive goblet cell numbers showed no change. mRNA expression in iNOS, eNOS, and nNOS showed no change. Irinotecan-induced mucositis is associated with altered secretory regulatory compounds; this may affect mucin secretion increasing the severity of irinotecan induced mucositis.

## Introduction

Alimentary mucositis (AM) a side effect of chemotherapy and radiotherapy occurs in 40% of standard dose chemotherapy patients, affecting the entire alimentary tract from mouth to anus ^1–5^. Symptoms can include one or multiple of mouth ulceration, abdominal pain, nausea, vomiting, abdominal bloating and diarrhoea ^2, 6, 7^. AM often leads to early cessation of treatment, which decreased the chance of remission and survival ^4, 8^. AM also increases the need for nutritional adjuncts, including fluid replacement, liquid diets, and total parenteral nutrition. The additional supportive care measures required for patients translates to a substantial cost increase per cycle of chemotherapy ^4, 5^. Currently, there is no effective prevention for AM ^9–14^, and treatment is limited to the management of symptoms once they occur, which is not always effective.

Mucus protects the mucosa from mechanical, chemical, and biological stress, therefore regulation of mucus secretion may be important during mucositis ^15–17^. Changes in mucus composition and secretion during mucositis have been demonstrated in numerous studies ^17–23^. These changes may be due to regulatory compounds such as nitric oxide (NO), vasoactive intestinal polypeptide (VIP) and prostaglandin E2 (PGE2), which may also be altered following chemotherapy.

Nitric Oxide (NO) is synthesised in the body through three isoforms: inducible nitric oxide synthase (iNOS), epithelial nitric oxide synthase (eNOS), and neuronal nitric oxide synthase (nNOS). NO has been previously shown to be associated with mucositis. A 5-Flurouracil (5-FU) oral mucositis model in hamsters showed a significant (p<0.05) increase in NO 10 days after 5-FU administration ^24^. With a significant (p<0.05) reduction in 5-FU induced oral damage when iNOS was selectively inhibited, but no significant reduction in oral damage when a non-selective NOS inhibitor was administered ^24^. These findings suggest that iNOS signalling may play a key role in oral mucositis, and eNOS and nNOS signalling may conversely contribute to maintaining gastrointestinal homeostasis.

VIP is associated with the regulation of goblet cell number and function. VIP increases goblet cell secretion ^25–27^, with VIP (100nM, 0.25 mL/min) perfusion significantly (p<0.05) increase cavitated cells and mucin secretion compared to controls after 30 min of perfusion ^27^. While the VIP receptor (VPAC1 and VPAC2) agonist [D-p- ClPhe6,Leu17]-VIP has been shown to decrease by 50% (p<0.01) goblet cell counts in the crypts and villi compared to vehicle and decrease 5-Ethynyl2’-deoxyuridine positive cells an indicator of cell proliferation by 77% (p<0.01) in a C57BL/6 mouse organotypic intestinal slice model. ^28^. This suggests VIP may decrease proliferation of goblet cells, while increasing mucin secretion. Mucositis studies in rat models have also shown an increase in mucus secretion from goblet cells and a decrease in goblet cell number ^17, 20, 21, 23, 29^, suggesting VIP may play a role in regulating mucin secretion in the development of chemotherapy-induced mucositis.

PGE2 have been shown to increase mucin secretion and cavitated goblet cells in the colon compared to controls ^27, 30,31^. 16,16’-dimethyl-PGE2 (dmPGE2) (2.5µM) perfusion significantly (p<0.05) increase cavitated cells and mucin secretion compared to controls after 30 min of perfusion in an infused isolated vascularly perfused rat colon model ^27^. Administration of PGE2 stimulated mucin secretion in LS174T human colonic adenocarcinoma cells in a dose dependent manner ^30^ and administration of dmPGE2 on the human colonic adenocarcinoma derived mucus-secreting goblet cell line (HT29-18N2) accelerated secretion of mucin glycoproteins ^31^. Furthermore, goblet cells in the GI tract expression mRNA for PGE2 specific prostaglandin receptors EP_1_ EP2 EP_3_ and EP_4_ in tissue from Sprague– Dawley rats ^32^. Highlighting the importance of PGE2 in mucin secretion across species. In mucositis studies using rat models of mucositis there has been a significant increase cavitation in goblet cells, a marker for mucin secretion ^17, 20, 21, 23, 29^, suggesting PGE2 may play a role in regulating mucin secretion in the development of chemotherapy-induced mucositis.

The aim of this study is to investigate regulator factors involved in mucin secretion (VIP, NO and PGE2) in the gastrointestinal tract, suspected of being involved in mucositis. In the irinotecan induced tumour bearing DA rat model of mucositis.

## Methods

### Ethics

Approval for the experiments was granted by the University of Adelaide and the Institute of Medical and Veterinary Science Animal Ethics Committees (approval number M-2010-118A). The conducted experiments fully comply with the National Health and Medical Research Council; Australia Code for the care and use of animals for scientific purposes (2013).

### Experimental design

The Dark Agouti Mammary Adenocarcinoma model used in this study has been extensively utilized and outlines in previous studies ^33–36^. Female Dark Agouti rats between 150g and 170g housed in a day night cycle 14:10 h at 22 ± 1°C were tumour inoculated via subcutaneous (s.c.) injection with 4.0×106 cells in 0.5 mL sterile PBS into each flank, 10 days prior to irinotecan administration. Rats were monitored four times a day and euthanized if they had a dull ruffled coat with accompanying dull and sunken eyes, were cold to touch with no spontaneous movement and a hunched appearance or had a tumour size greater than 10% of its body weight.

Rats were randomly assigned to control (6), 6 h (5), 24 h (5), 48 h (5), 72 h (8), 96 h (5), 120 h (6) as previously described ^17, 23^ Atropine was administered 0.01 mg/kg s.c. reducing the cholinergic reaction from irinotecan.

Irinotecan 175 mg/kg (i.p.) was administered in sorbitol/lactic acid buffer (45 mg/ml sorbitol, 0.9 mg/ml lactic acid, pH 3.4) (supplied by Pfizer, Kalamazoo, Michigan, USA). The control group was administered atropine and sorbitol/lactic acid buffer only. Rats were killed with 3% isofluorane in 100% O_2_, exsanguination and then cervical dislocation.

The gastrointestinal tract (GIT) was dissected as previously described ^17, 23^. Briefly, the alimentary tract was dissected out from the pyloric sphincter to the rectum The small intestine (SI) and colon was flushed with chilled, sterile distilled water. Dissected small and large intestine samples were fixed in 10% neutral buffered formalin, processed, and embedded in paraffin for histology. Dissected small intestine mucosa was scrapped and collected, then stored at -80°C for RNA extraction.

### Immunohistochemistry

To investigate whether iNOS, eNOS, VIP and PGE2 expression is affected by chemotherapy, anti-iNOS, anti-eNOS, anti-VIP and anti-PGE2 antibodies (Abcam, Cambridge, UK) were used following the method previously described ^17, 23^. Sections were cut at 4μm, deparaffinised in xylene and rehydrated in a graded serious of ethanol’s. Antigen retrieval with 10 mM citrate buffer (pH 6.0) occurred. Followed by blocking endogenous peroxidases with 3% hydrogen peroxide in methanol, of non-specific binding with blocking solution (Ultra Streptavidin HRP kit, Signet, Dedham, Massachusetts, U.S.A), and of endogenous avidin and biotin with an Avidin Biotin kit (Vector Laboratories, Burlingame, California, U.S.A.).

Sections were incubated at room temperature for 1 h with rabbit polyclonal antibodies: anti-iNOS (1:100 dilution, 2.0 µg/ml), anti-eNOS (1:20 dilution, 25.0 µg/ml), anti-VIP (1:300 dilution), or anti-PGE2 antibody (1:200 dilution) (Abcam, Cambridge, UK). Then incubated at room temperature with linking reagent (Ultra Streptavidin HRP kit for 30 min, and Labelling reagent (Ultra Streptavidin HRP kit) for 30 min. Sections were then stained with 3,3’- diaminobenzidine (DAB) and counterstained using Haematoxylin. Dehydrating through a graded series of ethanol then cover slipped. Staining intensity (0 = negative; 1 = weak; 2 = moderate; 3 = strong; 4 = very intense) was used to analyse the change in expression based on a previously validated technique ^37^.

### RNA isolation and reverse transcription

RNA was extracted from mucosal scraping using Aurum™ Total RNA Mini Kit (Bio-Rad, California, USA) and converted to cDNA with iScript™ cDNA Synthesis Kit (Bio-Rad, California, USA) as previously described ^23^. Then diluted to a working concentration of 100 ng/µL.

### Real Time PCR

Gene expression of iNOS and eNOS was investigated using Real-time PCR, Rotor-Gene Q (Qiagen, Sydney, Australia). The amplification mixes contained 0.5 µL each of forward and reverse primers, 5 µL of 2x SYBR green (Bio-Rad Laboratories Inc. USA), 3 µL nuclease free water and 1 µL of cDNA from the mucosal samples. Thermal cycling conditions for each primer pair were optimized (Supplementary Table 1). Two housekeeping genes Ubiquitin C (UBC) and beta-2-microglobulin (B2M) in conjunction with Pfaffle quantification were used to determine relative gene expression ^38^.

### Statistical analysis

Statistical analysis with Kruskal-Wallis test with a secondary Dunn’s comparative test or a Mann Whitney U test were performed, using GraphPad Prism 6.0 software (Graphpad software, Califorina, U.S.A). Further analysis with non-parametric regressions were also performed using StataIC12 (StataCorp LP, Texas, USA). The differences between mean ranks were determined to be significant at p < 0.05. Cohen’s D tests were used to determine effect size with the effect considered small if d > 0.20, moderate if d > 0.50, and large if d > 0.80 ^39^.

## Results

### iNOS expression

#### iNOS stained goblet cells

The mean staining intensity of iNOS positive goblet cells of vehicle control rats in the jejunum was 2.0 ± 0.8 (mean ± SEM) in the villi and 1.4 ± 0.9 in the crypts, and 2.3 ± 0.6 in the colon. Following irinotecan administration, the staining intensity tended to decrease, and was lowest at 48 h in the jejunum (0.8 ± 0.5 and 1.0 ± 0.6 for villus and crypt, respectively), and 96 h in the colon (1.0 ± 0.7) (Supplementary Figure 1).

The mean number of goblets cells positively stained with iNOS in the jejunum of vehicle control rats was 1.6 ± 0.2 per villus and 0.7 ± 0.2 per crypt. Following irinotecan administration, the mean number of cells positively stained with iNOS was lowest in the villi at 48 h (0.3 ± 0.2 per villus, d = 2.89, large effect) and significantly decreased in the crypts at 48 h (0.1 ± 0.07 per crypt, p<0.05, d = 2.42, large effect) (Figure 1).

**Figure 1.**
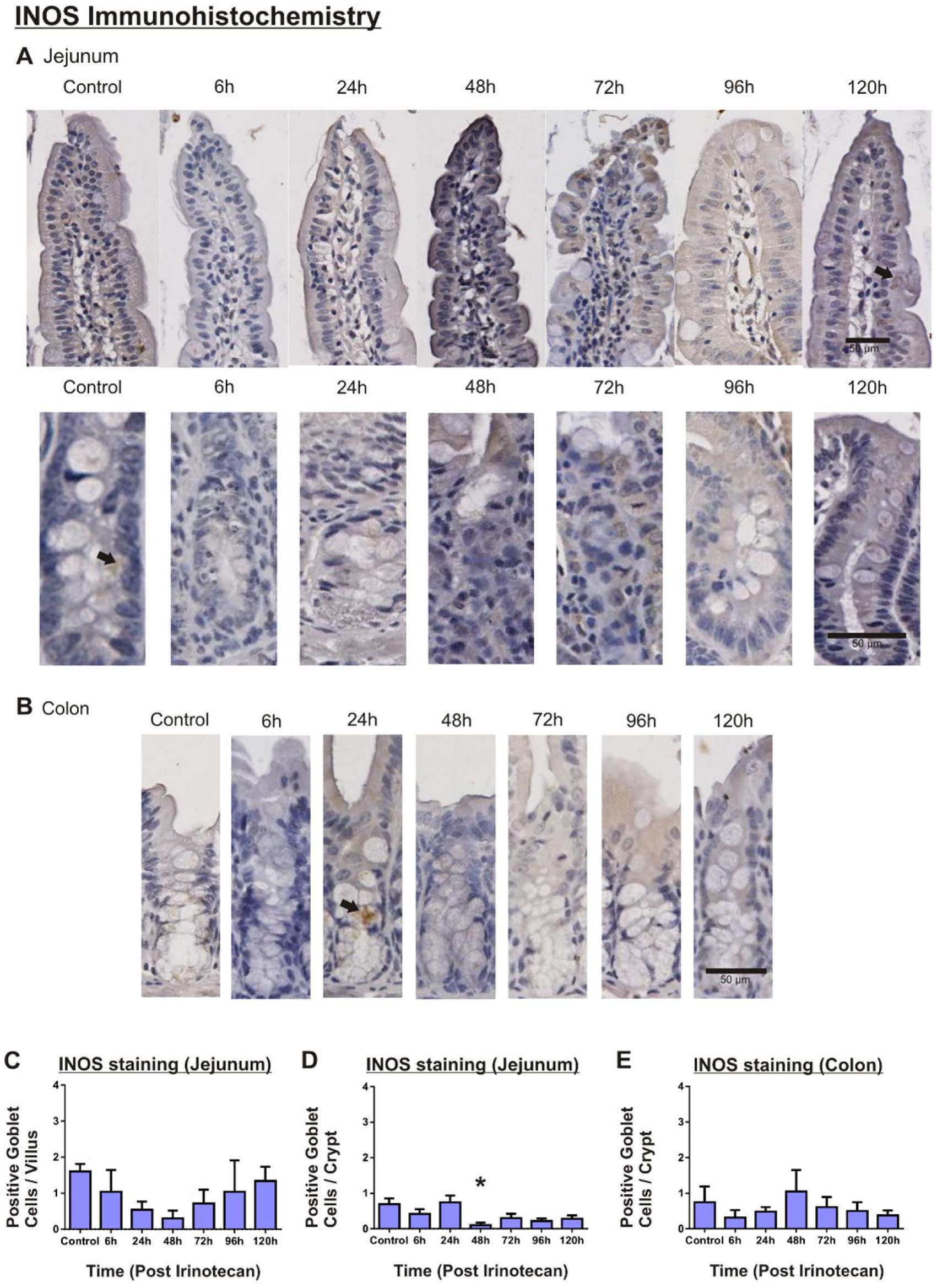
Immunohistochemistry of iNOS positively stained goblet cells. A. Jejunum (villus and crypt). Arrow, iNOS positive goblet cell. B. Colon. Arrow, iNOS positively stained goblet cell (original magnification 20X). C-E. iNOS positive goblet cell counts in the jejunum villus (C) and crypts (D), and colon (E) (* denotes statistical significance, where p<0.05).

The mean number of goblet cells positively stained with iNOS in the colon of vehicle control rats was 0.75 ± 0.44 per crypt. Following irinotecan administration, the mean number of positively stained cells was lowest at 6 h (0.3 ± 0.6 per crypt, d = 0.55, medium effect), and highest at 48 h (1.1 ± 0.6 per crypt, d = 0.26, small effect), although not significant (Figure 1).

#### iNOS stained enteric ganglia

The mean staining intensity of iNOS positive stained enteric ganglia in the myenteric plexus of vehicle control rats was 0.5 ± 0.3 in the jejunum, and 1.5 ± 0.7 in the colon. Following irinotecan administration, the staining intensity was lowest in the jejunum at 24 h (0.0 ± 0.0, d = 0.37, small effect) and lowest in the colon at 96 h (0.5 ± 0.3, d = 1.49, large effect) (Supplementary Figure 1).

The mean number of enteric ganglia positively stained with iNOS in the myenteric plexus of control rats was 2.3 ± 0.8 per mm in the jejunum, and 1.0 ± 0.7 per mm in the colon. The mean number of iNOS-positive enteric ganglia in the myenteric plexus varied following irinotecan administration, being lowest at 24 h in the jejunum (1.3 ± 0.8 per mm, d = 0.61, medium effect), and highest at 48 h (4.0 ± 1.1 per mm, d = 0.79, medium effect). The mean number of iNOS-positive enteric ganglia in the myenteric plexus of the colon tended to increase, and was highest at 120 h in the colon (4.8 ± 0.8 per mm, d = 2.36, large effect) (Supplementary Figure 2).

#### iNOS gene expression

In the jejunum, iNOS gene expression in the vehicle control group was 1.6 ± 0.3. The relative gene expression (fold change) was highest at 72 h (2.6 ±1.1) following irinotecan compared to control. However, there was variability within groups, resulting in mean ranks not being significantly different (Figure 2).

**Figure 2.**
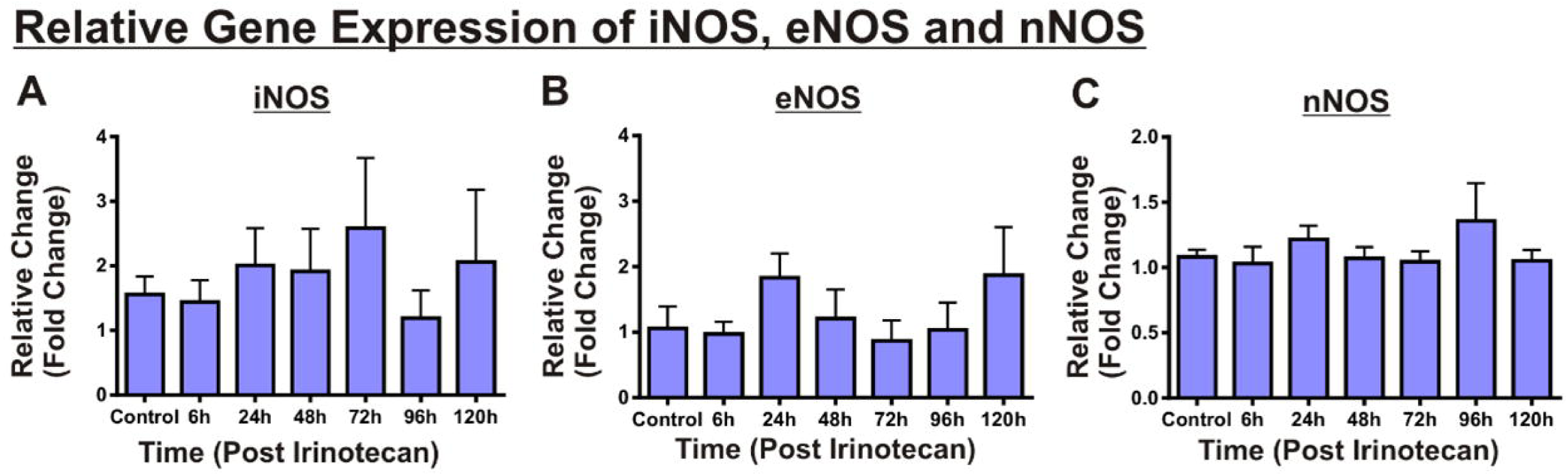
Expression of iNOS, eNOS and nNOS in the jejunum of irinotecan treated rats compared to vehicle controls. A. iNOS, B. eNOS, and C. nNOS following irinotecan administration.

### eNOS expression

#### eNOS stained goblet cells

The mean staining intensity of eNOS positive goblet cells of vehicle control rats was 0.8 ± 0.4 in the villi and 1.0 ± 0.4 in the crypts, and 3.2 ± 0.4 in the colon. Following irinotecan administration, the staining intensity was highest at 120 h in the villi (2.2 ± 0.2) and 6 h (1.8 ± 0.6) in the crypts. Staining intensity was lower in the colon following irinotecan, being lowest at 96 h (1.0 ± 0.6) (Supplementary Figure 1).

The mean number of goblets cells positively stained with eNOS in the jejunum of vehicle control rats was 2.3 ± 0.5, and 0.6 ± 0.2 per villus and crypt, respectively. Following irinotecan administration, the mean number of cells positively stained with eNOS tended to be lower than control and was lowest in the villi at 96 h (0.7 ± 0.2 per villus, d = 1.88, large effect) and in the crypts at 48 h (0.1 ± 0.1 per crypt, d = 1.73, large effect) (Figure 3).

**Figure 3.**
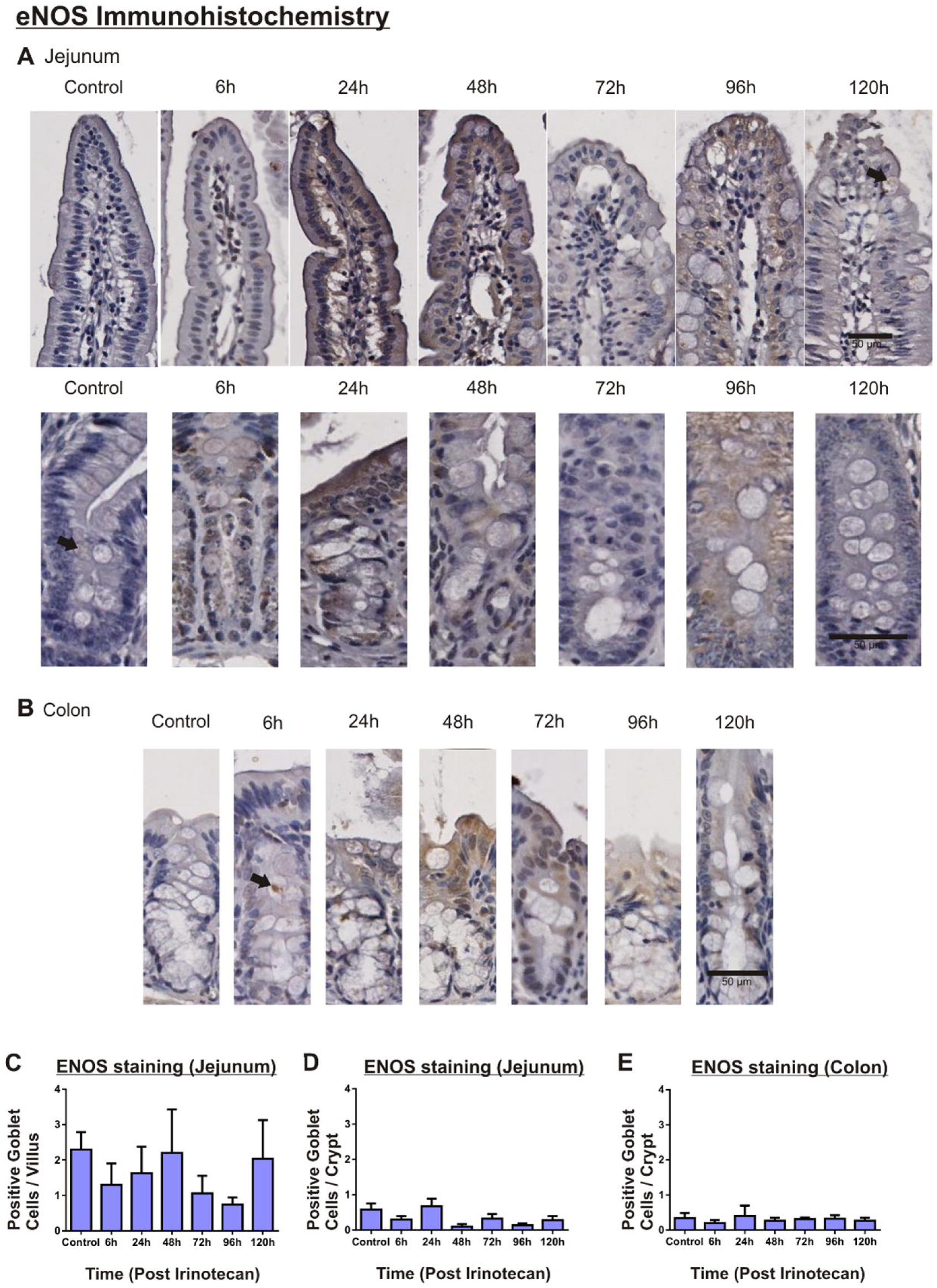
Immunohistochemistry of eNOS positively stained goblet cells. A. Jejunum, villus and crypt sections. Arrow, eNOS positive goblet cell B. Colon, crypt sections. Arrow, eNOS positive goblet cell. C. Jejunum, counts of eNOS positive goblet cells per villus. D. Jejunum, counts of eNOS positive goblet cells per crypt E. Colon, counts of eNOS positive goblet cells per crypt. (Original magnification 20X).

The mean number of goblet cells positively stained with eNOS in the colon of vehicle control rats was 0.34 ± 0.15 per crypt. Following irinotecan administration, the mean number of positively stained cells did not change (Figure 3).

#### eNOS stained enteric ganglia

The mean staining intensity of eNOS positive enteric ganglia in the myenteric plexus of vehicle control rats was 0.5 ± 0.3 in the jejunum, and 1.5 ± 0.7 in the colon. Following irinotecan administration, the staining intensity was variable, increasing and decreasing at different time points. Staining intensity was lowest in the jejunum at 24 h (0.0 ± 0.0, d = 1.20, large effect) and highest at 48 h (2.3 ± 0.9, d = 1.38, large effect). Intensity was lowest in the colon at 48 h (0.5 ± 0.3 per mm, d = 0.90, large effect) and highest in the colon at 120 h (2.3 ± 0.8, d = 0.48, small effect) (Supplementary Figure 1).

The mean number of eNOS positive enteric ganglia in the myenteric plexus of vehicle control rats was 0.7 ± 0.4 per mm in the jejunum, and 1.5 ± 0.8 per mm in the colon. Following irinotecan administration, the number of positively stained enteric ganglia in the jejunum followed a similar variable pattern to the staining intensity. The number of eNOS positive enteric ganglia was lowest in the jejunum at 24 h (0.0 ± 0.0 per mm, d = 1.29, large effect), and highest at 48 h (2.8 ± 1.3 per mm, d = 1.18, large effect). The mean number of eNOS positive enteric ganglia was highest in the colon at 120 h (2.8 ± 0.6 per mm, d = 0.75, large effect), although not significant (Supplementary Figure 2).

#### eNOS gene expression

In the jejunum, eNOS gene expression in vehicle controls was 1.1 ± 0.3. Following irinotecan administration, the highest gene expression occurred at 24 h (1.8 ± 0.4 fold change) and 120 h (1.9 ± 0.7 fold change) compared to controls. However, this did not reach significance (Figure 2).

### nNOS

#### nNOS gene expression

In the jejunum, nNOS mRNA expression was 1.1 ± 0.1 in vehicle control rats. No change in expression was observed in irinotecan treated rats compared to controls (Figure 2).

### VIP expression

#### VIP stained goblet cells

The mean staining intensity of VIP positive goblet cells in the jejunum of vehicle control rats was 1.2 ± 0.2 and 1.5 ± 0.2 in the villi and crypts, respectively, and 1.8 ± 0.4 in the colon. Following irinotecan administration, the staining intensity did not change significantly in the villi or crypts of the jejunum, although varied from as high as 2.5 ± 0.5 (48 h) to as low as 1.0 ± 0.0 (120 h) in the colon (Supplementary Figure 3).

The number of goblets cells positively stained for VIP in the jejunum of control rats was 2.8 ± 0.6 per villus and 2.1 ± 0.5 per crypt. Following irinotecan administration, the number of positively stained cells varied, being lowest in the villi at 6 h (1.7 ± 0.6 per villus, d = 0.83, large effect), and then highest at 96 h (6.3 ± 2.0 per villus, d = 1.2, large effect). The number of VIP positive cells in the crypts also varied following irinotecan administration, being lowest at 48 h (0.9 ± 0.5 per crypt, d = 1.11, large effect), and highest at 120 h (3.0 ± 0.8 per crypt, d = 0.58, medium effect) (Figure 4).

**Figure 4.**
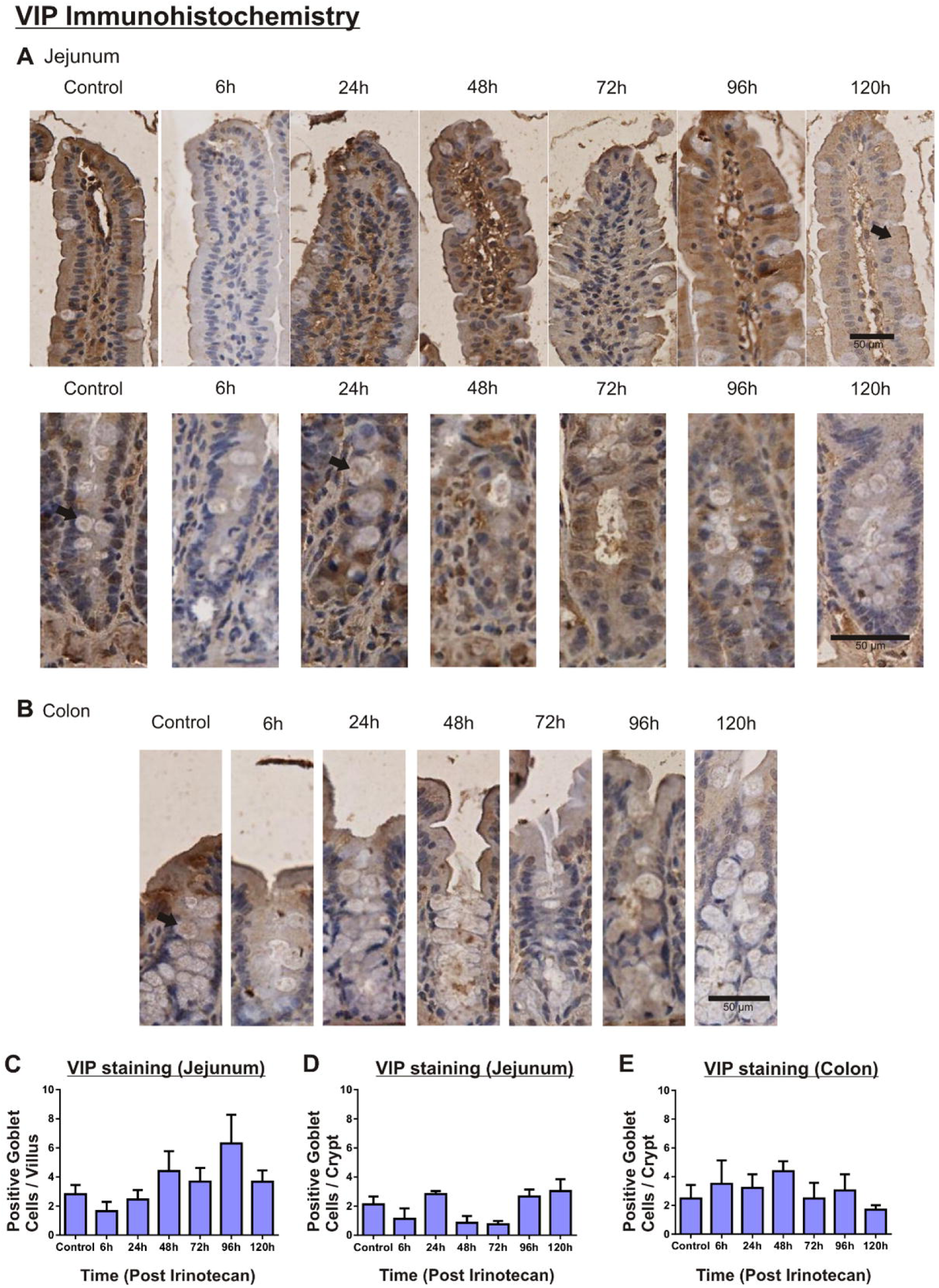
Immunohistochemistry of VIP positively stained goblet cells. A. Jejunum, villus and crypt sections. Arrow, VIP positively stained goblet cell. B. Colon, crypt sections. Arrows, VIP positively stained goblet cell C. Jejunum, counts of VIP positive goblet cells per villus. D. Jejunum, counts of VIP positive goblet cells per crypt E. Colon, counts of VIP positive goblet cells per crypt. (Original Magnification 20X).

The number of goblet cells positively stained for VIP in the colon of control rats was 2.5 ± 0.9 per crypt. Following irinotecan administration the number of positively stained cells varied, being highest at 48 h (4.4 ± 0.7 per crypt, d = 1.03, large effect), and lowest at 120 h (1.7 ± 0.3 per crypt, d = 0.52, medium effect) (Figure 4).

#### VIP stained enteric ganglia

The number of VIP positive enteric ganglia in vehicle control rats was 0.8 ± 0.3 per mm in the jejunum, and 1.2 ± 0.7 per mm in the colon. Following irinotecan administration the number of positively stained enteric ganglia was lowest in the jejunum at 6 h (0.4 ± 0.6 per mm, d = 0.67, medium effect) and was highest at 48 h (2.2 ± 1.0 per mm, d = 0.90, large effect). In the colon, the number of VIP positive enteric ganglia was highest at 24 h (2.6 ± 0.5 per mm, d = 1.04, large effect), and lowest at 48 h (0.6 ± 0.4 pre mm, d = 0.45, small effect) (Supplementary Figure 2).

### PGE2 expression

#### PGE2 stained goblet cells

The mean staining intensity of PGE2 positive goblet cells of control rats in the jejunum was 2.0 ± 0.3 in the villi and 2.0 ± 0.4 in the crypts, and 1.1 ± 0.4 in the colon. Following irinotecan administration staining intensity in the goblet cells was lowest in the jejunum at 6 and 24 h (1.5 ± 0.3 and 1.5 ± 0.3 in the villi and crypts, respectively), and highest at 120 h (2.3 ± 0.6) in both the villi and crypts). Intensity was highest in the colon at 24 h (2.3 ± 0.4), and lowest at controls (Supplementary Figure 3).

The number of goblets cells positively stained with PGE2 in the jejunum of vehicle control rats were 5.1 ± 1.3 per villus, and 2.2 ± 0.5 per crypt. Following irinotecan administration, the number of cells positively stained was lowest in the villi at 6 h (3.4 ± 1.0 per villus, d = 0.69, medium effect) and was highest at 24 h (7.6 ± 2.5 per villus, d = 0.61, large effect). The number of cells positively stained was highest in the crypts at 24h (4.6 ± 0.7, d = 1.74, large effect), and then lowest at 72 h (0.6 ± 0.3 per crypt, d = 1.54, large effect) (Figure 5).

**Figure 5.**
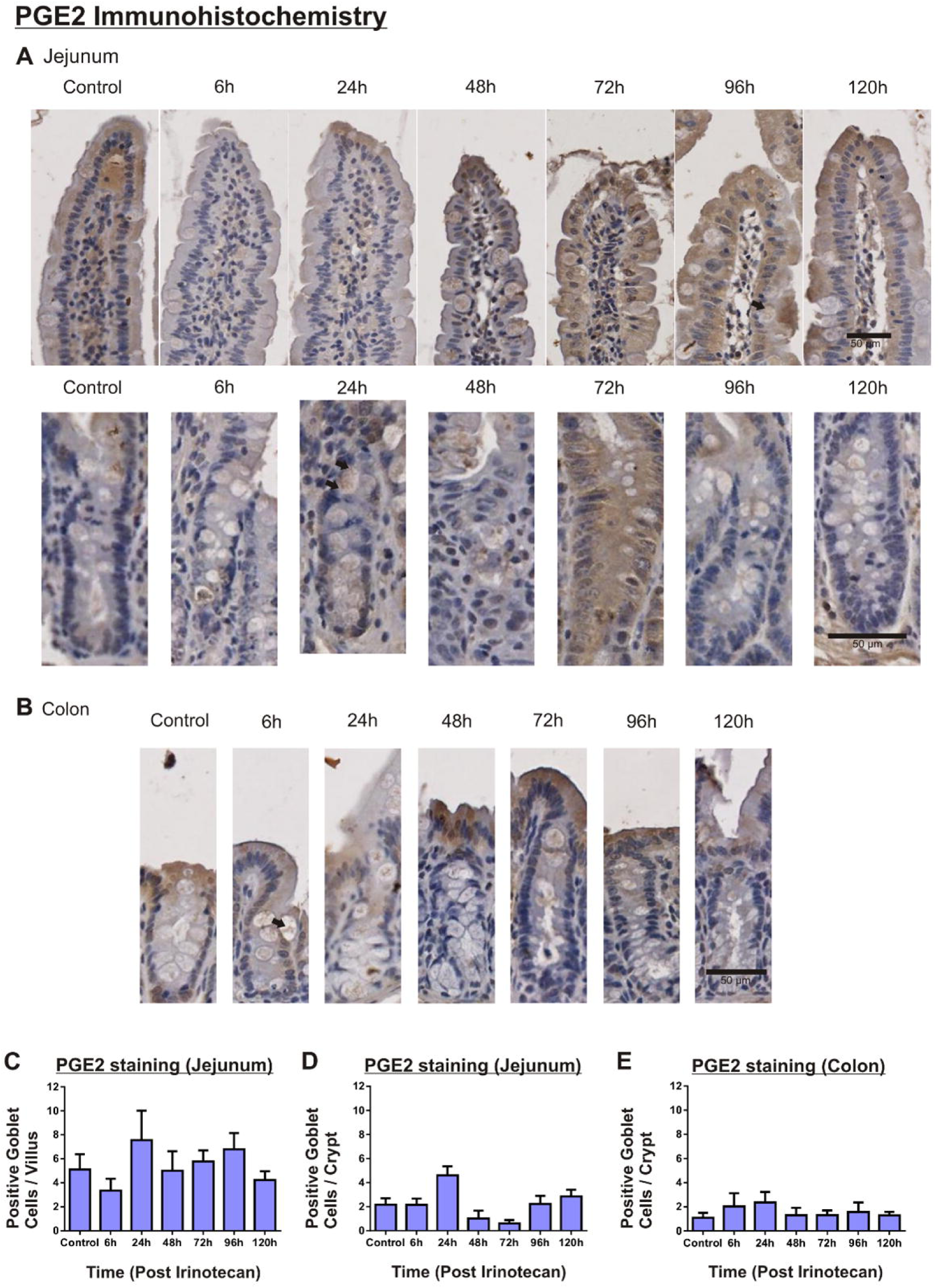
Immunohistochemistry of PGE2 positively stained goblet cells. A. Jejunum, villus and crypt sections. B. Colon, crypt sections C. Jejunum, counts of PGE2 positive goblet cells per villus. D. Jejunum, counts of PGE2 positive goblet cells per crypt E. Colon, counts of PGE2 positive goblet cells per crypt. (Original Magnification 20X).

The number of goblet cells positively stained with PGE2 in the colon of vehicle control rats was 1.1 ± 0.4 per crypt. Following irinotecan administration, the number of cells positively stained with PGE2 was highest in the crypts at 24 h (2.3 ± 0.9 per crypt, d = 0.88, large effect), and lowest at 120 h (1.3 ± 0.3 per crypt, d = 0.24, small effect) (Figure 5).

#### PGE2 stained enteric ganglia

The number of enteric ganglia positively stained for PGE2 in vehicle control rats in the jejunum was 1.3 ± 1.0 per mm, and in the colon was 0.6 ± 0.4 per mm. Following irinotecan administration, the number of PGE2 positively stained enteric ganglia in the jejunum was lowest at 24 h (0.8 ± 0.4 per mm, d = 0.34, small effect) and highest at 48 h (3.0 ± 1.5 per mm, d = 0.67, medium effect). In the colon, the number of PGE2 positive enteric ganglia tended to increase, being highest at 24 h (2.6 ± 0.5 per mm, d = 1.97, large effect) (Supplementary Figure 2).

### Non-parametric regression

Non-parametric regression analyses were carried out for parameters of the jejunum and colon. In the jejunum, strong correlations were demonstrated between iNOS gene expression, eNOS gene expression and nNOS gene expression. Furthermore, strong correlations were demonstrated between VIP and PGE2 positive goblet cell counts in the villi, as well as VIP and PGE2, VIP and iNOS, PGE2 and iNOS, and iNOS and eNOS positive goblet cells in the crypts. Strong correlations were also demonstrated between PGE2 and iNOS, iNOS and eNOS, and VIP and PGE2 positive enteric ganglia in the myenteric plexus (Supplementary Table 2) (Figure 6).

**Figure 6.**
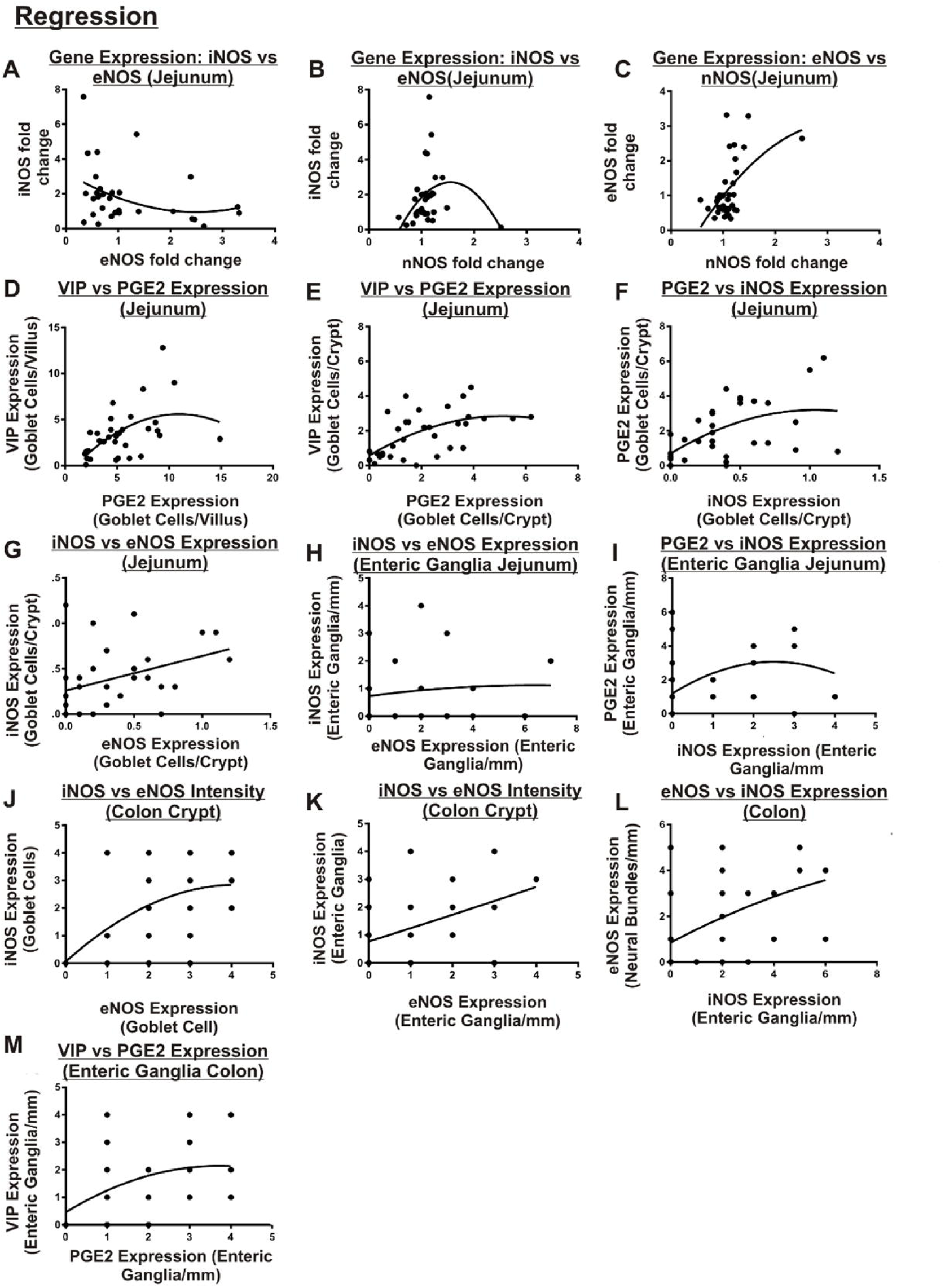
Non parametric regressions analysis of iNOS, eNOS and nNOS expression, VIP, PGE2, iNOS and eNOS positive cells and PGE2, iNOSeNOS and VIP positive enteric ganglia. A-C. Gene expression (jejunum): iNOS vs eNOS (A), iNOS vs nNOS (B), eNOS vs nNOS (C). D-I. IHC expression (jejunum): VIP vs PGE2 positive cells/villus (D), VIP vs PGE2 positive cells/crypt (E), PGE2 vs iNOS positive cells/crypt (F), iNOS vs eNOS positive cells/crypt (G), iNOS vs eNOS positive Enteric Ganglia/mm (H), PGE2 vs iNOS positive Enteric Ganglia/mm (I). J-M IHC expression (colon): iNOS vs eNOS intensity (crypt) (J), iNOS vs eNOS intensity (Enteric Ganglia) (K), eNOS vs iNOS positive Enteric Ganglia/mm (L), VIP vs PGE2 positive Enteric Ganglia/mm (M).

In the colon, strong correlations were demonstrated between iNOS and eNOS staining intensity of goblet cells, iNOS and eNOS staining intensity of enteric ganglia, and staining intensity of enteric ganglia between eNOS and iNOS, and VIP and PGE2 (Supplementary Table 2) (Figure 6).

## Discussion

GI mucositis is a severe, dose-limiting, toxic side effect of cancer treatment ^4, 5^. This study has demonstrated for the first time that neurotransmitters and regulatory chemicals thought to be involved in secretion (particularly in goblet cells) are modified during mucositis. These chemicals and neurotransmitters may therefore affect the rate and amount of mucus secretion from goblet cells, impacting on the protective barrier of the intestinal mucosa. Previous studies have demonstrated intestinal goblet cells and mucins to be significantly affected by chemotherapy agents, 5- FU and irinotecan ^19, 20^. However, these studies have used non-tumour bearing rats, and did not investigate the regulatory chemicals and neurotransmitters that may be involved in this process.

This study demonstrates decreased iNOS positive goblet cells in the jejunum (significant in the crypts). Gene expression of iNOS did not change significantly over the time course of mucositis, suggesting the decrease in iNOS positive cells is post-translational and not due to down-regulation of genetic expression. The findings of this study differ from those of a previous study using an irinotecan-induced mouse model of mucositis (once a day for 4 days of 60 mg/kg i.p., C57Bl/6), which showed a significant increase in iNOS protein in the duodenum 24h after concluding treatment during mucositis using western blots ^40^. This discrepancy may be due to the different dosing schedule (4x daily smaller doses vs 1x larger single dose), or regional differences between the duodenum and jejunum caused by earlier exposure to SN-38 (converted from SN-38G as its delivered to the duodenum via the bile duct), and highlights the need for further investigation into the role of iNOS in mucositis.

Non-parametric regression analyses showed correlations between iNOS and eNOS gene expression, and iNOS and nNOS gene expression, suggesting limited change to iNOS, eNOS and nNOS gene regulation. iNOS and eNOS positive goblet cells were correlated in the jejunum crypts, as were iNOS and eNOS positive enteric ganglia in the myenteric plexus of the jejunum, and iNOS and eNOS goblet cell intensity, and myenteric plexus enteric ganglia intensity and positive enteric ganglia/mm. These findings suggest that iNOS and eNOS protein expression is similar in the jejunum. iNOS is Ca²⁺ independent and eNOS is Ca²⁺ dependent ^41, 42^, suggesting that Ca²⁺ levels in the jejunum may not be altered during mucositis. This however does not mean that signalling and activity exerted by Ca²⁺ are not altered during mucositis. Again, further investigation into this phenomenon is warranted.

VIP expression tended to increase following treatment with irinotecan, with positive enteric ganglia highest in the myenteric plexus at 48 h in the jejunum, and at 6 and 24 h in the colon, and VIP positive goblet cells at the highest level in the villi of the jejunum at 48 h. VIP stimulates secretory activity ^27, 43, 44^ by activation of two G-protein-coupled receptors, VPAC1 and VPAC2 ^45^ in the presence of extracellular Ca^2+^ ^46^. Increased VIP expression may therefore lead to increased mucin secretion during mucositis. This is consistent with previous studies in rat models of chemotherapy-induced mucositis ^17, 20, 21, 23, 29^, which have demonstrated an increase in mucin secretion by goblet cells following irinotecan administration.

PGE2 positive enteric ganglia were highest in the myenteric plexus of the jejunum at 48 h, and were elevated in the colon in irinotecan treated rats, although not significant. PGE2 positive goblet cells were also increased in the jejunum and colon. This may suggest that cyclooxygenase (COX)-2 and arachidonic acid may also be increased during mucositis, as PGE2 is the product of the metallisation of arachidonic acid by COX-2. Previous studies have shown COX-2 expression to increase significantly (p<0.001) in submucosal tissues in response to targeted radiation in a hamster buccal pouch model of mucositis ^47^. Mucin secretion may also be increased in response to increases in PGE2, as previous studies have demonstrated associations between PGE2 and increased mucin secretion ^27, 31^, offering a potential explanation for the increased mucin secretion observed in previous studies ^19, 20^.

Correlations between VIP and PGE2 positive enteric ganglia in the myenteric plexus, and goblet cells of the jejunum demonstrated in this study were an interesting finding. Karaki and Kuwahara (2004) suggested that increased levels of PGE2 can either directly increase secretion, or act upon VIP secretomotor neurons to increase secretion in the intestine ^48^. The findings of this study suggest this link between PGE2 and VIP may be a potential mechanism for the increased secretion from intestinal goblet cells. Therefore, further investigations into the specific actions of PGE2 and VIP during mucositis is warranted.

In conclusion, iNOS expression is decreased following irinotecan. Regulatory compounds and enteric neurotransmitters may be involved with the control of secretion and expression of mucins. These compounds may regulate ENS signalling, and as a result are likely to be involved in the pathophysiology of mucositis. Further studies are warranted to elucidate the role of these compounds on ENS signalling and regulation of mucin secretion in the development of chemotherapy-induced mucositis.

## Acknowledgments

This study was funded by National Health and Medical Research Council funding (1016696)

---

Supplementary Table 1. Primer sequence and cycling conditions for real time PCR, UBC, B2M, iNOS, and eNOS.

Supplementary Table 2. Non-parametric regression analyses of parameters in the jejunum and colon.

**<u>Supplementary 1</u>** Immunohistochemistry (staining intensity). A. iNOS: Jejunum goblet cells (villus). B. iNOS: Jejunum goblet cells (crypt). C. iNOS: Colon goblet cells. D. iNOS: Jejunum myenteric plexus. E. iNOS: Colon myenteric plexus. F. eNOS: Jejunum goblet cells (villus). G. eNOS: Jejunum goblet cells (crypt). H. eNOS: Colon goblet cells. I. eNOS: Jejunum myenteric plexus. J. eNOS: Colon myenteric plexus.

**<u>Supplementary 2</u>** Immunohistochemistry: counts of iNOS, eNOS, VIP and PGE2 positively stained enteric ganglia (myenteric plexus). A. iNOS: Jejunum, enteric ganglia per mm. B. iNOS: Colon, enteric ganglia per mm. C. eNOS: Jejunum, enteric ganglia per mm. D. eNOS: Colon, enteric ganglia per mm. E. VIP: Jejunum, enteric ganglia per mm. F. VIP: Colon, enteric ganglia per mm. G. PGE2: Jejunum, enteric ganglia per mm. H. PGE2: Colon, enteric ganglia per mm. (* denotes statistical significance, where p<0.05).

**<u>Supplementary 3</u>** Immunohistochemistry staining intensity of positively stained cells. A. VIP: Jejunum, villus goblet cells. B. VIP: Jejunum, crypt goblet cells. C. VIP: Colon, goblet cells. D. PGE2: Jejunum, villus goblet cells. E. PGE2: Jejunum, crypt goblet cells. F. PGE2: Colon, goblet cells.

## References

1. Keefe DM, Brealey J, Goland GJ, et al. Chemotherapy for cancer causes apoptosis that precedes hypoplasia in crypts of the small intestine in humans. Gut 2000; 47: 632–637. 2000/10/18.

2. Keefe DM, Cummins AG, Dale BM, et al. Effect of high-dose chemotherapy on intestinal permeability in humans. Clin Sci (Lond) 1997; 92: 385–389. 1997/04/01.

3. Pico JL, Avila-Garavito A and Naccache P. Mucositis: Its Occurrence, Consequences, and Treatment in the Oncology Setting. Oncologist 1998; 3: 446–451. 1999/07/01.

4. Elting LS, Cooksley C, Chambers M, et al. The burdens of cancer therapy. Clinical and economic outcomes of chemotherapy-induced mucositis. Cancer 2003; 98: 1531–1539. 2003/09/26. DOI: 10.1002/cncr.11671.

5. Elting LS, Cooksley CD, Chambers MS, et al. Risk, outcomes, and costs of radiation-induced oral mucositis among patients with head-and-neck malignancies. Int J Radiat Oncol Biol Phys 2007; 68: 1110–1120. 2007/04/03. DOI: 10.1016/j.ijrobp.2007.01.053.

6. Keefe DM. Gastrointestinal mucositis: a new biological model. Support Care Cancer 2004; 12: 6–9. 2003/11/08. DOI: 10.1007/s00520-003-0550-9.

7. Sonis ST, Costa JW, Jr., Evitts SM, et al. Effect of epidermal growth factor on ulcerative mucositis in hamsters that receive cancer chemotherapy. Oral Surg Oral Med Oral Pathol 1992; 74: 749–755. 1992/12/11.

8. Savarese DM, Hsieh C and Stewart FM. Clinical impact of chemotherapy dose escalation in patients with hematologic malignancies and solid tumors. J Clin Oncol 1997; 15: 2981–2995. 1997/08/01.

9. Keefe DM, Schubert MM, Elting LS, et al. Updated clinical practice guidelines for the prevention and treatment of mucositis. Cancer 2007; 109: 820–831. 2007/01/20. DOI: 10.1002/cncr.22484.

10. Sonis ST, Elting LS, Keefe D, et al. Perspectives on cancer therapy-induced mucosal injury: pathogenesis, measurement, epidemiology, and consequences for patients. Cancer 2004; 100: 1995–2025. 2004/04/27. DOI: 10.1002/cncr.20162.

11. Rubenstein EB, Peterson DE, Schubert M, et al. Clinical practice guidelines for the prevention and treatment of cancer therapy-induced oral and gastrointestinal mucositis. Cancer 2004; 100: 2026–2046. 2004/04/27. DOI: 10.1002/cncr.20163.

12. Saltz L, Shimada Y and Khayat D. CPT-11 (irinotecan) and 5-fluorouracil: a promising combination for therapy of colorectal cancer. Eur J Cancer 1996; 32A Suppl 3: S24–31. 1996/01/01.

13. Saltz LB. Understanding and managing chemotherapy-induced diarrhea. J Support Oncol 2003; 1: 35-46; discussion 38-41, 45–36. 2004/09/09.

14. Wadler S, Benson AB, 3rd, Engelking C, et al. Recommended guidelines for the treatment of chemotherapy-induced diarrhea. J Clin Oncol 1998; 16: 3169–3178. 1998/09/17.

15. Thorpe D, Stringer A and Butler R. Chemotherapy-induced mucositis: the role of mucin secretion and regulation, and the enteric nervous system. Neurotoxicology 2013; 38: 101–105. 2013/07/06. DOI: 10.1016/j.neuro.2013.06.007.

16. Thorpe D. The role of mucins in mucositis. Current opinion in supportive and palliative care 2019; 13: 114–118. 2019/03/21. DOI: 10.1097/spc.0000000000000423.

17. Thorpe D, Butler R, Sultani M, et al. Irinotecan-Induced Mucositis Is Associated with Goblet Cell Dysregulation and Neural Cell Damage in a Tumour Bearing DA Rat Model. Pathology oncology research : POR 2019 2019/03/29. DOI: 10.1007/s12253-019-00644-x.

18. de Koning BA, Sluis M, Lindenbergh-Kortleve DJ, et al. Methotrexate-induced mucositis in mucin 2-deficient mice. J Cell Physiol 2007; 210: 144–152. 2006/09/26. DOI: 10.1002/jcp.20822.

19. Stringer AM, Gibson RJ, Logan RM, et al. Irinotecan-induced mucositis is associated with changes in intestinal mucins. Cancer Chemother Pharmacol 2009; 64: 123–132. 2008/11/11. DOI: 10.1007/s00280-008-0855-y.

20. Stringer AM, Gibson RJ, Logan RM, et al. Gastrointestinal microflora and mucins may play a critical role in the development of 5-Fluorouracil-induced gastrointestinal mucositis. Exp Biol Med (Maywood) 2009; 234: 430–441. 2009/01/30. DOI: 10.3181/0810-rm-301.

21. Saegusa Y, Ichikawa T, Iwai T, et al. Changes in the mucus barrier of the rat during 5-fluorouracil-induced gastrointestinal mucositis. Scand J Gastroenterol 2008; 43: 59–65. 2008/10/22.

22. Verburg M, Renes IB, Meijer HP, et al. Selective sparing of goblet cells and paneth cells in the intestine of methotrexate-treated rats. Am J Physiol Gastrointest Liver Physiol 2000; 279: G1037–1047. 2000/10/29.

23. Thorpe D, Sultani M and Stringer A. Irinotecan induces enterocyte cell death and changes to muc2 and muc4 composition during mucositis in a tumour-bearing DA rat model. Cancer Chemother Pharmacol 2019; 83: 893–904. 2019/03/01. DOI: 10.1007/s00280-019-03787-5.

24. Leitao RF, Ribeiro RA, Bellaguarda EA, et al. Role of nitric oxide on pathogenesis of 5-fluorouracil induced experimental oral mucositis in hamster. Cancer Chemother Pharmacol 2007; 59: 603–612. 2006/09/01. DOI: 10.1007/s00280-006-0301-y.

25. Kirkegaard P, Lundberg JM, Poulsen SS, et al. Vasoactive intestinal polypeptidergic nerves and Brunner’s gland secretion in the rat. Gastroenterology 1981; 81: 872–878. 1981/11/01.

26. Dartt DA, Kessler TL, Chung E-H, et al. Vasoactive Intestinal Peptide-Stimulated Glycoconjugate Secretion from Conjunctival Goblet Cells. Experimental Eye Research 1996; 63: 27–33. DOI: 10.1006/exer.1996.0088.

27. Plaisancie P, Barcelo A, Moro F, et al. Effects of neurotransmitters, gut hormones, and inflammatory mediators on mucus discharge in rat colon. Am J Physiol 1998; 275: G1073–1084. 1998/11/14.

28. Schwerdtfeger LA and Tobet SA. Vasoactive intestinal peptide regulates ileal goblet cell production in mice. Physiological Reports 2020; 8. Article. DOI: 10.14814/phy2.14363.

29. Stringer AM, Gibson RJ, Bowen JM, et al. Irinotecan-induced mucositis manifesting as diarrhoea corresponds with an amended intestinal flora and mucin profile. Int J Exp Pathol 2009; 90: 489–499. 2009/09/22. DOI: 10.1111/j.1365-2613.2009.00671.x.

30. Wright DH, Ford-Hutchinson AW, Chadee K, et al. The human prostanoid DP receptor stimulates mucin secretion in LS174T cells. British Journal of Pharmacology 2000; 131: 1537–1545. Article. DOI: 10.1038/sj.bjp.0703688.

31. Phillips TE, Stanley CM and Wilson J. The effect of 16,16-dimethyl prostaglandin E2 on proliferation of an intestinal goblet cell line and its synthesis and secretion of mucin glycoproteins. Prostaglandins Leukot Essent Fatty Acids 1993; 48: 423–428. 1993/06/01.

32. Northey A, Denis D, Cirino M, et al. Cellular distribution of prostanoid EP receptors mRNA in the rat gastrointestinal tract. Prostaglandins & Other Lipid Mediators 2000; 62: 145–156. DOI: 10.1016/S0090-6980(00)00058-7.

33. Gibson RJ, Keefe DM, Thompson FM, et al. Effect of interleukin-11 on ameliorating intestinal damage after methotrexate treatment of breast cancer in rats. Dig Dis Sci 2002; 47: 2751–2757. 2002/12/25.

34. Gibson RJ, Keefe DM, Clarke JM, et al. The effect of keratinocyte growth factor on tumour growth and small intestinal mucositis after chemotherapy in the rat with breast cancer. Cancer Chemother Pharmacol 2002; 50: 53–58. 2002/07/12. DOI: 10.1007/s00280-002-0460-4.

35. Bowen JM, Gibson RJ, Cummins AG, et al. Irinotecan changes gene expression in the small intestine of the rat with breast cancer. Cancer Chemother Pharmacol 2007; 59: 337–348. 2006/06/27. DOI: 10.1007/s00280-006-0275-9.

36. Gibson RJ, Bowen JM and Keefe DM. Palifermin reduces diarrhea and increases survival following irinotecan treatment in tumor-bearing DA rats. Int J Cancer 2005; 116: 464–470. 2005/04/01. DOI: 10.1002/ijc.21082.

37. Bowen JM, Gibson RJ, Keefe DM, et al. Cytotoxic chemotherapy upregulates pro-apoptotic Bax and Bak in the small intestine of rats and humans. Pathology 2005; 37: 56–62. 2005/05/07.

38. Pfaffl MW. Quantification strategies in real-time PCR. In: Bustin S (ed) A-Z of quantitative PCR. La Jolla, CA, USA: International University Line, 2004, pp.87-112.

39. Cohen J. Statistical Power Analysis for the Behavioral Sciences. Taylor & Francis, 2013.

40. Lima-Junior RC, Figueiredo AA, Freitas HC, et al. Involvement of nitric oxide on the pathogenesis of irinotecan-induced intestinal mucositis: role of cytokines on inducible nitric oxide synthase activation. Cancer Chemother Pharmacol 2012; 69: 931–942. 2011/11/22. DOI: 10.1007/s00280-011-1780-z.

41. Nathan C. Inducible nitric oxide synthase: what difference does it make? J Clin Invest 1997; 100: 2417–2423. 1997/11/20. DOI: 10.1172/jci119782.

42. Wallace JL and Miller MJ. Nitric oxide in mucosal defense: a little goes a long way. Gastroenterology 2000; 119: 512–520. 2000/08/10.

43. Dickson L and Finlayson K. VPAC and PAC receptors: From ligands to function. Pharmacol Ther 2009; 121: 294–316. 2008/12/27. DOI: 10.1016/j.pharmthera.2008.11.006.

44. Laburthe M, Augeron C, Rouyer-Fessard C, et al. Functional VIP receptors in the human mucus-secreting colonic epithelial cell line CL.16E. Am J Physiol 1989; 256: G443-450. 1989/03/01.

45. Martin B, Lopez de Maturana R, Brenneman R, et al. Class II G protein-coupled receptors and their ligands in neuronal function and protection. Neuromolecular Med 2005; 7: 3–36. 2005/07/30.

46. Bou-Hanna C, Berthon B, Combettes L, et al. Role of calcium in carbachol- and neurotensin-induced mucin exocytosis in a human colonic goblet cell line and cross-talk with the cyclic AMP pathway. Biochem J 1994; 299 ( Pt 2): 579–585. 1994/04/15.

47. Sonis ST, O’Donnell KE, Popat R, et al. The relationship between mucosal cyclooxygenase-2 (COX-2) expression and experimental radiation-induced mucositis. Oral Oncol 2004; 40: 170–176. 2003/12/25.

48. Karaki SI and Kuwahara A. Regulation of intestinal secretion involved in the interaction between neurotransmitters and prostaglandin E2. Neurogastroenterology and motility : the official journal of the European Gastrointestinal Motility Society 2004; 16 Suppl 1: 96–99. 2004/04/07. DOI: 10.1111/j.1743-3150.2004.00482.x.

