## Supplementary Table 1 for "Irinotecan-induced mucositis is associated with variation in the expression of regulatory compounds associated with goblet cells"

**Table 1** Primer sequence and cycling conditions of real time PCR.

| **Gene** | **Primer sequence (5’ - 3’)** | **Amplicon length** | **Annealing (˚C)** | **Cycles** |
| --- | --- | --- | --- | --- |
| **UBC** | F:TCGTACCTTTCTCACCACAGTATCTAG  R: GAAAACTAAGACACCTCCCCATCA | 82 |  |  |
| **B2M** | F: CGAGACCGATGTATATGCTTGC  R: GTCCAGATGATTCAGAGCTCCA | 114 |  |  |
| **iNOS** | F:GGGAGCCAGAGCAGTACAAG  R:CATGGTGAACACGTTCTTGG | 95 | 50 | 50 |
| **eNOS** | F: GCTTGGGATCCCTGGTATTT  R: GCTTGACCCAATAGCTGCTC | 85 | 50 | 50 |
