## Supplementary Table 2 for "Irinotecan-induced mucositis is associated with variation in the expression of regulatory compounds associated with goblet cells"

**Supplementary Data Table 2** Significant (p<0.05) non-parametric regressions.

| **Gut Region** | **Dependent Variable** | **Independent Variable** | **Coefficient** | **95% CI Coefficient** | **Pseudo R²** |
| --- | --- | --- | --- | --- | --- |
| **Jejunum** | iNOS Gene Expression | eNOS Gene Expression | -0.61 | -1.21  -0.00 | 0.10 |
| **Jejunum** | iNOS Gene Expression | nNOS Gene Expression | 0.73 | 0.52  3.51 | 0.01 |
| **Jejunum** | eNOS Gene Expression | nNOS Gene Expression | 1.23 | 0.24  2.22 | 0.13 |
| **Jejunum** | eNOS Gene Expression | eNOS IHC intensity (Enteric Ganglia) | 0.46 | -0.00  0.93 | 0.06 |
| **Jejunum** | nNOS Gene Expression | eNOS IHC Expression (Cells/Crypt) | -0.25 | -0.49  -0.01 | 0.05 |
| **Jejunum** | nNOS Gene Expression | PGE2 IHC intensity (Crypt) | 0.10 | 0.00  0.20 | 0.08 |
| **Jejunum** | VIP IHC Expression (Cells/Villus) | PGE2 IHC Expression (Cells/Villus) | 0.42 | 0.06  0.78 | 0.09 |
| **Jejunum** | VIP IHC Expression (Cells/Crypt) | PGE2 IHC Expression (Cells/Crypt) | 0.46 | 0.13  0.80 | 0.26 |
| **Jejunum** | VIP IHC Expression (Cells/Crypt) | iNOS IHC Intensity (Crypts) | 0.85 | 0.26  1.44 | 0.06 |
| **Jejunum** | PGE2 IHC Expression (Cells/Crypt) | iNOS IHC Expression (Cells/Crypt) | 4.25 | 1.59  6.91 | 0.11 |
| **Jejunum** | PGE2 IHC Expression (Cells/Crypt) | VIP IHC Expression (Cells/Crypt) | -1.25 | -2.20  0.30 | 0.17 |
| **Jejunum** | VIP IHC Expression (Enteric Ganglia per mm) | VIP IHC Intensity (Crypt) | 1.00 | 0.42  1.58 | 0.15 |
| **Jejunum** | PGE2 IHC Expression (Enteric Ganglia per mm) | iNOS IHC Expression (Enteric Ganglia per mm) | 1.33 | 0.70  1.96 | 0.19 |
| **Jejunum** | iNOS IHC Expression (Villus) | iNOS IHC Expression (Enteric Ganglia per mm) | 0.38 | 0.01  0.74 | 0.12 |
| **Jejunum** | iNOS IHC Expression (Villus) | iNOS IHC Intensity (Villus) | 0.45 | 0.12  0.78 | 0.13 |
| **Jejunum** | iNOS IHC Expression (Villus) | eNOS IHC Intensity (Villus) | 0.50 | 0.01  0.99 | 0.05 |
| **Jejunum** | iNOS IHC Expression (Crypt) | eNOS IHC Expression (Crypt) | 0.40 | 0.03  0.78 | 0.11 |
| **Jejunum** | iNOS IHC Expression (Crypt) | iNOS IHC Intensity (Crypt) | 0.20 | 0.11  0.29 | 0.15 |
| **Jejunum** | iNOS IHC Expression (Enteric Ganglia per mm) | eNOS IHC Expression (Enteric Ganglia per mm) | 0.25 | 0.03  0.47 | 0.10 |
| **Jejunum** | eNOS IHC Expression (Enteric Ganglia per mm) | eNOS IHC intensity (Enteric Ganglia) | 1.00 | 0.68  1.32 | 0.32 |
| **Jejunum** | eNOS IHC intensity (Enteric Ganglia) | eNOS IHC intensity (Villus) | 0.33 | 0.01  0.66 | 0.039 |
| **Jejunum** | iNOS IHC Expression (Crypt) | iNOS IHC intensity (Crypt) | 1.00 | 0.52  1.48 | 0.03 |
| **Jejunum** | eNOS IHC intensity (Villus) | eNOS IHC intensity (Crypt) | 1.00 | 0.63  1.37 | 0.24 |
| **Jejunum** | PGE2 IHC intensity (Villus) | PGE2 IHC intensity (Crypt) | 0.50 | 0.26  0.74 | 0.09 |
| **Colon** | VIP IHC Expression (Enteric Ganglia) | PGE2 IHC Expression (Enteric Ganglia per mm) | 0.67 | 0.22  1.11 | 0.13 |
| **Colon** | eNOS IHC Expression (Enteric Ganglia per mm) | iNOS IHC Expression (Enteric Ganglia per mm) | 0.75 | 0.35  1.15 | 0.27 |
| **Colon** | eNOS IHC Expression (Enteric Ganglia per mm) | iNOS IHC intensity (Enteric Ganglia) | 1.00 | 0.33  1.67 | 0.22 |
| **Colon** | eNOS IHC Expression (Enteric Ganglia per mm) | eNOS IHC intensity (Enteric Ganglia) | 1.00 | 0.40  1.60 | 0.33 |
| **Colon** | iNOS IHC Expression (Enteric Ganglia per mm) | iNOS IHC intensity (Enteric Ganglia) | 1.25 | 0.87  1.63 | 0.41 |
| **Colon** | iNOS IHC Expression (Enteric Ganglia per mm) | eNOS IHC intensity (Enteric Ganglia) | 1.00 | 0.17  1.83 | 0.06 |
| **Colon** | iNOS IHC Intensity (Enteric Ganglia) | eNOS IHC intensity (Enteric Ganglia) | 0.67 | 0.15  1.19 | 0.20 |
| **Colon** | iNOS IHC Intensity (Crypt) | eNOS IHC intensity (Crypt) | 1.00 | 0.54  1.46 | 0.31 |
