## Supplementary figures and images for "Irinotecan-induced mucositis is associated with variation in the expression of regulatory compounds associated with goblet cells"

### Supplementary Figure 1

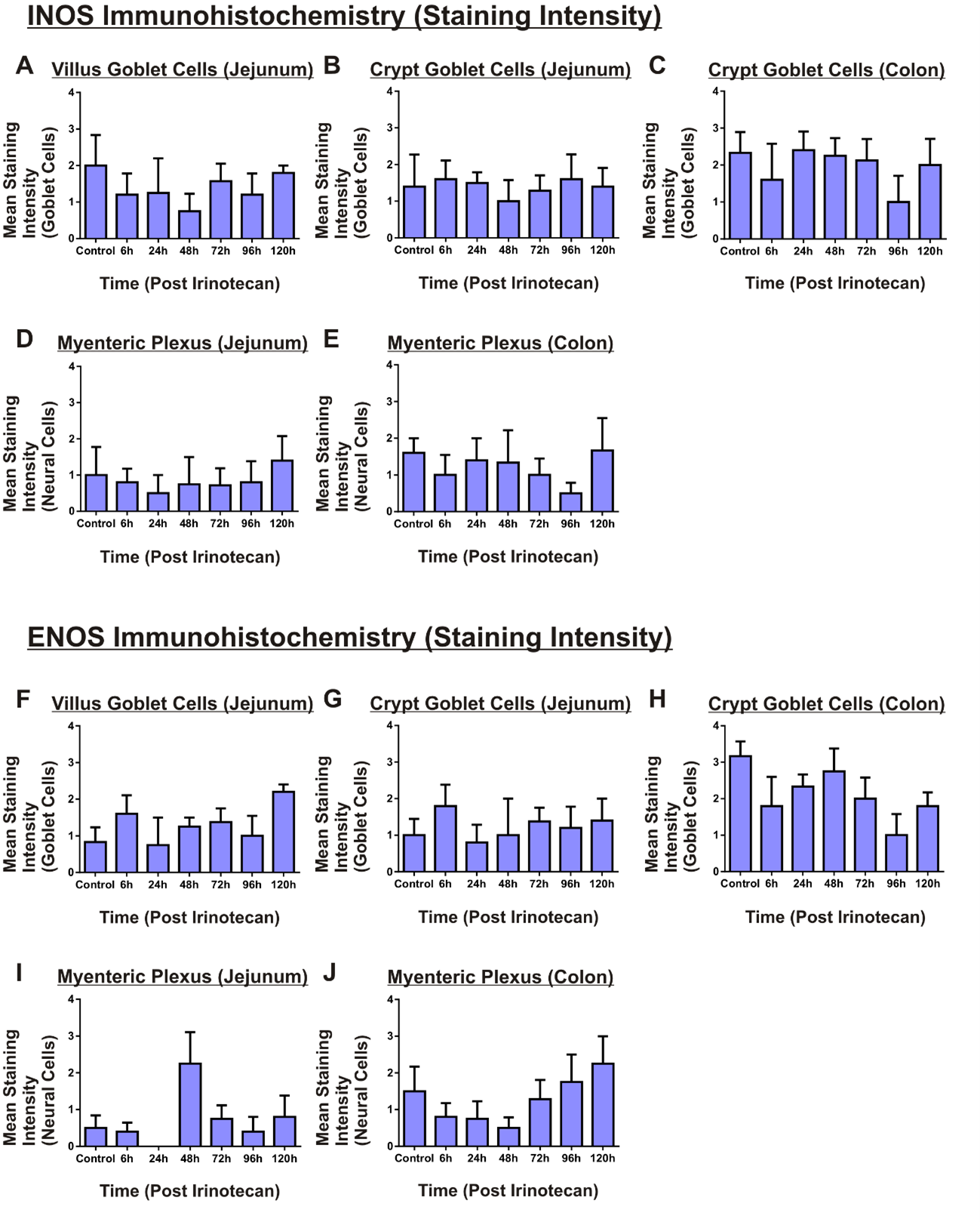

### Supplementary Figure 2

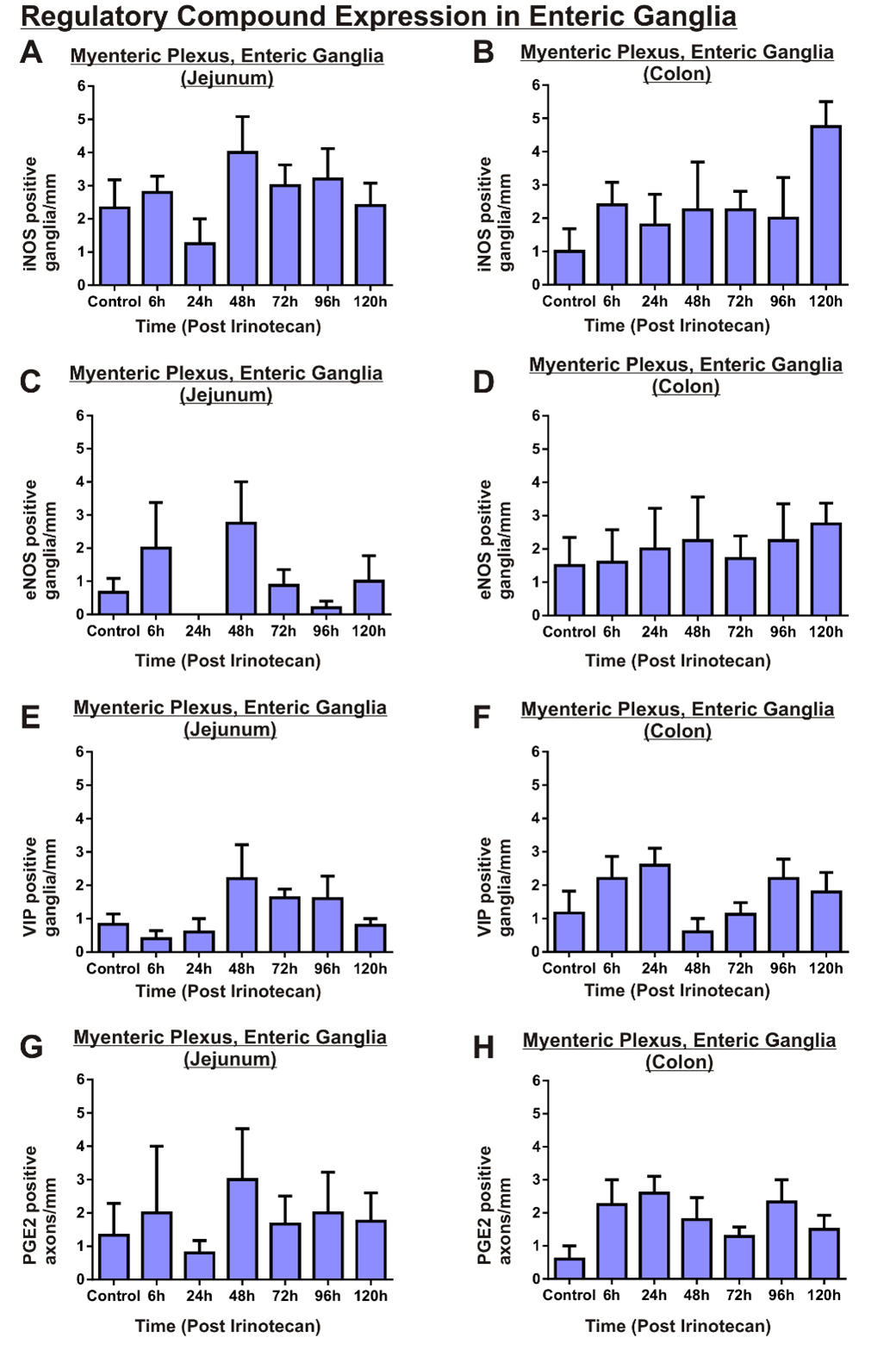

### Supplementary Figure 3

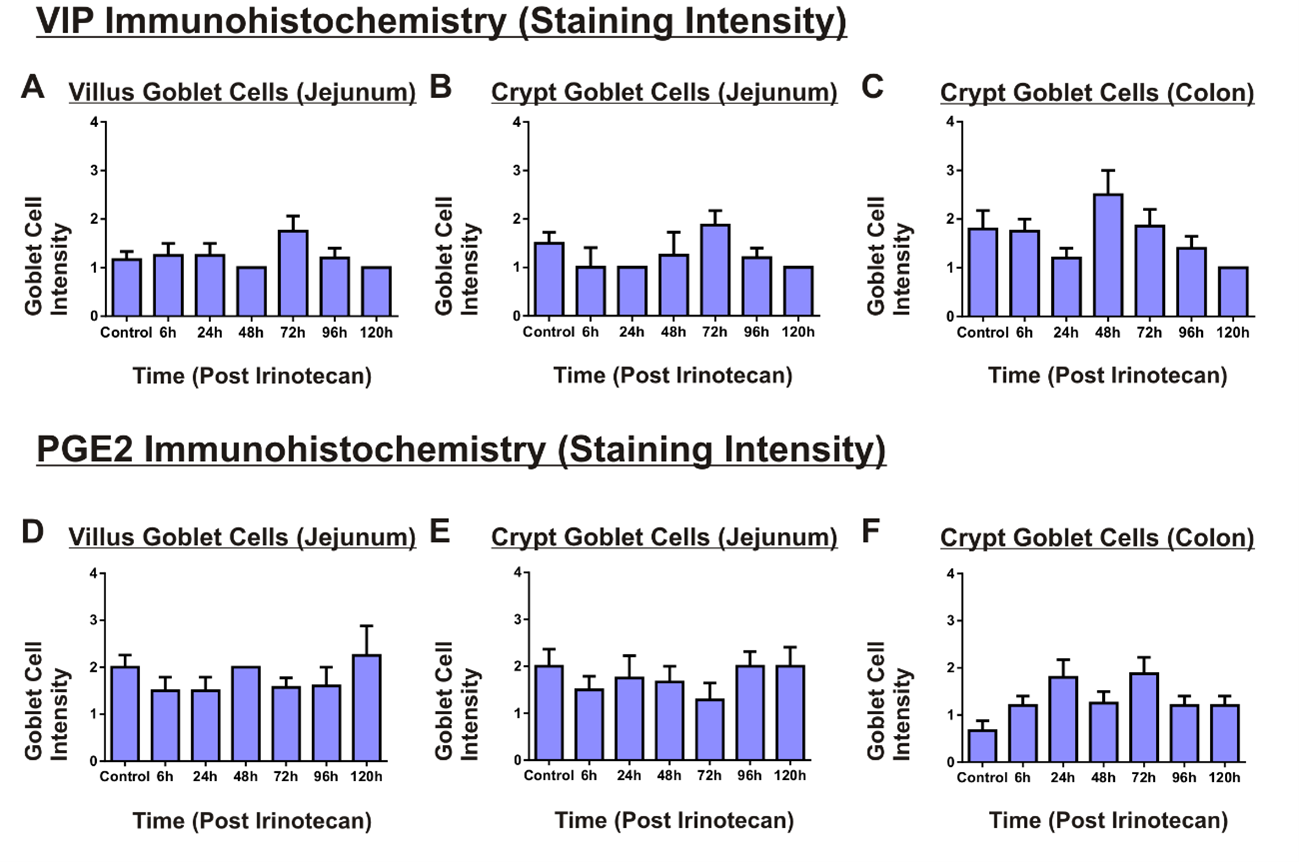
